# Spondweni virus causes fetal harm in a mouse model of vertical transmission and is transmitted by *Aedes aegypti* mosquitoes

**DOI:** 10.1101/824466

**Authors:** Anna S. Jaeger, Andrea M. Weiler, Ryan V. Moriarty, Sierra Rybarczyk, Shelby L. O’Connor, David H. O’Connor, Davis M. Seelig, Michael K. Fritsch, Thomas C. Friedrich, Matthew T. Aliota

## Abstract

Spondweni virus (SPONV) is the most closely related known flavivirus to Zika virus (ZIKV). Its pathogenic potential and vector specificity have not been well defined. SPONV has been found predominantly in Africa, but was recently detected in a pool of *Culex quinquefasciatus* mosquitoes in Haiti. Here we show that SPONV can cause significant fetal harm, including demise, comparable to ZIKV, in a mouse model of vertical transmission. Following maternal inoculation, we detected infectious SPONV in placentas and fetuses, along with significant fetal and placental histopathology, together indicating vertical transmission. To test vector competence, we exposed *Aedes aegypti* and *Culex quinquefasciatus* mosquitoes to SPONV-infected bloodmeals. *Aedes aegypti* could efficiently transmit SPONV, whereas *Culex quinquefasciatus* could not. Our results suggest that SPONV has the same features that made ZIKV a public health risk.

## Main

Zika virus (ZIKV) was originally isolated over seventy years ago, and was thought to cause a mild, self-limiting, febrile illness ^1,2^. Not until the outbreak in the Americas in 2015 and 2016 was ZIKV identified as a cause of significant adverse pregnancy outcomes ^3,4^. Before the definition of congenital Zika syndrome (CZS) in 2016, gestational arbovirus infection was not associated with birth defects. Spondweni virus (SPONV) is the closest known relative to ZIKV, but whether SPONV is an emerging threat to pregnant women and their babies is unknown. It was previously thought that SPONV was geographically confined to Africa and caused only mild disease in rare human infections, reminiscent of the consensus around ZIKV in the decades following its discovery, but recent data suggest that it is spreading beyond Africa ^5^. SPONV may therefore be poised to harm pregnancies in new, immunologically naive populations. To do this, SPONV would need to fulfill two major criteria: it would need to be vertically transmitted and cause fetal harm, and be transmitted between humans by the urban mosquito vector *Aedes aegypti*, which is associated with large-scale outbreaks of related arboviruses.

The first identification of SPONV was thought to have occurred in 1955 in South Africa ^6,7^. However, it was later recognized that SPONV was in fact isolated three years earlier in Nigeria, but was misidentified at the time as a strain of ZIKV because of serological cross-reactivity ^2,8,9^. Serological cross-reactivity with ZIKV and other flaviviruses likely still confounds accurate diagnostics today. As a result, only six well-documented clinical cases of SPONV infection have ever been described ^9–11^. It is likely that many infections have gone unrecognized—serosurveys have detected evidence of SPONV infection in 10 countries throughout Sub-Saharan Africa ^6,12–15^. Still, these six cases indicate that infection with SPONV typically involves an acute, self-limiting mild to moderate febrile illness ^6^, but a subset (4/6) of cases are believed to progress to more serious disease, including vascular leakage and neurological involvement ^6,9,10,16,17^; these symptoms resemble those of other infections common in the same regions ^6,8,18^. Nothing is known about the risks posed by SPONV infection during pregnancy in humans. Powassan virus and West Nile virus, ZIKV-related neurotropic flaviviruses, can infect human placental explants and cause fetal demise in mice ^19^. In addition, intrauterine infection with St. Louis encephalitis virus resulted in severe neurological outcomes in mice that was dependent on the gestational day of challenge ^20^. In pregnant mice whose type 1 interferon responses were blocked by antibody treatment, SPONV did not cause fetal harm, but the placenta and fetus were infected ^21^.

Potential vectors for SPONV have not been identified, although SPONV has been isolated from several mosquito genera ^17,22,23^. SPONV is considered to be exclusively mosquito-borne, but recent mouse experiments suggest that it can be sexually transmitted ^24^. Whether SPONV could spill over into a human-mosquito cycle involving *Aedes aegypti* is unclear. One recent study suggested strains of *Aedes albopictus*, *Aedes aegypti*, and *Culex quinquefasciatus* were all unable to transmit SPONV ^25^. Overall, with only limited studies evaluating SPONV’s pathogenic potential and vector specificity, there is little data to guide our expectations for the potential of SPONV to cause fetal harm and adapt to an urban or peri-urban transmission cycle involving *Aedes aegypti* and/or other human-biting mosquito species.

We aimed to better characterize both the pathogenic potential of SPONV during pregnancy and to also identify potential vectors of the virus. Using an established vertical transmission model in mice ^26^, we assessed fetal outcomes after infection at embryonic day 7.5 with SPONV as compared to both ZIKV and dengue virus (DENV). We found that SPONV caused fetal harm, similar to what is observed from ZIKV infection in this model. Vector competence experiments showed that *Ae. aegypti* could transmit SPONV when exposed to bloodmeal titers that approximate physiological titers, while *Cx. quinquefasciatus* could not. Our study contributes to the characterization of SPONV pathogenesis and identifies a potential urban vector for the virus, collectively suggesting that this seemingly esoteric virus has features that could portend medically significant future outbreaks.

## Results

### Type I interferon deficient mice are susceptible to SPONV infection

Recent studies have demonstrated that SPONV replicates in AG129 mice deficient in type I and II interferon ^24^ and in mice treated with an IFNAR1-blocking monoclonal antibody (mAb) ^21^. We sought to establish a model that was less immunocompromised than AG129 mice, and one in which transplacental ZIKV infection and fetal damage had been demonstrated ^26–28^ to better understand SPONV pathologic outcomes during pregnancy. First, groups of nonpregnant, mixed sex 6-to 11-week-old mice lacking type I interferon signaling (*Ifnar1*^−/−^) were inoculated subcutaneously (s.c.) in the footpad with 10^3^ or 10^2^ PFU of SPONV strain SA Ar94 (this is the only strain used in these studies, so it will be referred to simply as SPONV); or 10^2^ PFU of the highly pathogenic African-lineage ZIKV strain DAK AR 41524 (ZIKV-DAK) ^26^. SPONV strain SA Ar94 is 98.8% nucleotide identical with the SPONV genome recovered from mosquitoes in Haiti, and is the same strain that has been used in previous mouse studies ^21,24^. Serum was collected at 2, 4, and 6 days post-inoculation (dpi) to confirm infection and determine the replication kinetics of SPONV in nonpregnant *Ifnar1*^−/−^ mice. We also collected and tested serum at 7, 14, and 21 days from mice surviving SPONV inoculation, because sustained vRNA loads were observed with the IFNAR1-blocking mAb model ^21^. SPONV viral titer in the serum peaked at 4 dpi but was typically below the limit of detection at 2 dpi (Fig. 1a), and in surviving animals there was no evidence of sustained viremia. ZIKV-DAK viremia also peaked at 4 dpi and reached significantly higher titers at peak than either SPONV-inoculated group (one-way ANOVA with Tukey’s multiple comparisons test; SPONV 10^3^ vs. ZIKV-DAK: *p* <0.0001, SPONV 10^2^ vs. ZIKV-DAK: *p* = 0.0004).

**Figure 1.**
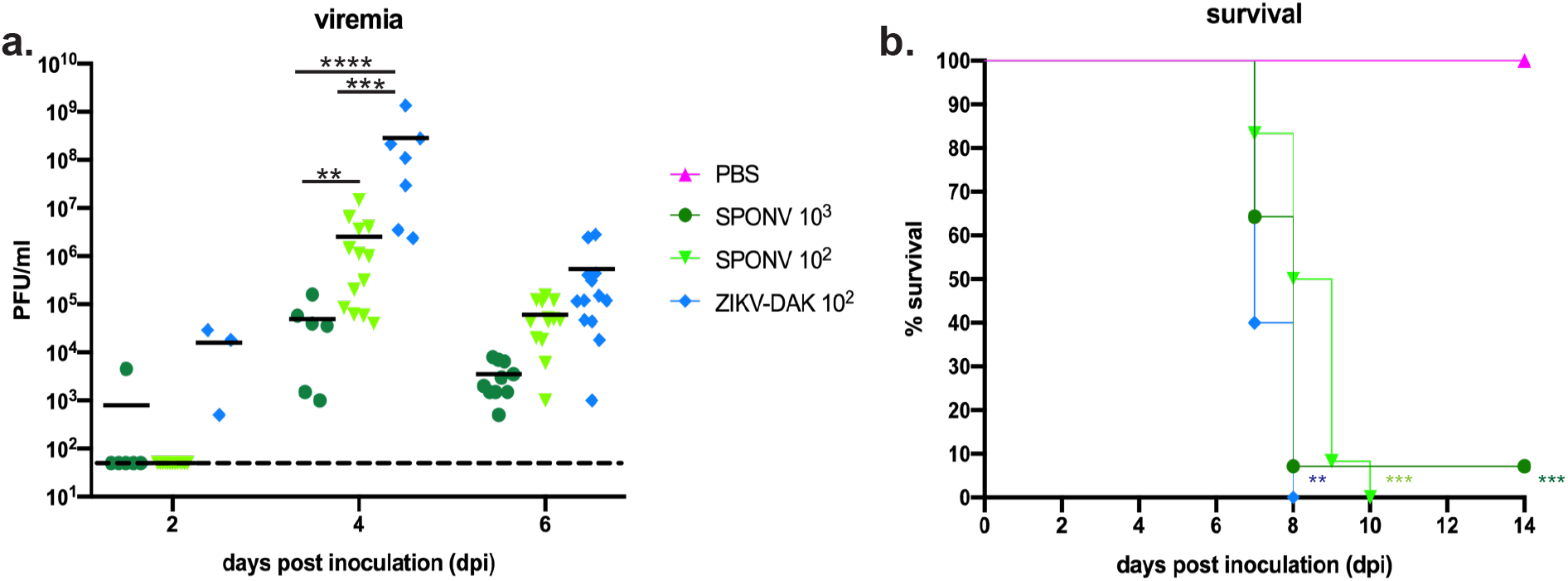
Characterization of SPONV in nonpregnant *Ifnar1*^−/−^ mice. **(a)** Serum was collected from mice during three independent replicates at 2, 4, and/or 6 days post inoculation and titered via plaque assay. The dotted line indicates the plaque assay limit of detection. Viremia peaked at 4dpi for all virus groups, with ZIKV-DAK replicating to significantly higher titers at 4dpi than SPONV (one-way ANOVA). \*\*\*\**p* < 0.0001; \*\*\**p* < 0.0002; \*\**p* < 0.002 **(b)** Survival curves of six-to eleven-week old *Ifnar1*^−/−^ mice subcutaneously inoculated with 10^3^ PFU of SPONV, 10^2^ PFU of SPONV, 10^2^ PFU ZIKV-DAK, or a PBS control. SPONV 10^3^: n=14; SPONV 10^2^: n=13, ZIKV-DAK: n=5; PBS: n=6). All virus infections caused significant mortality by 14dpi as compared to PBS controls (Fisher’s exact test). \*\*\**p* < 0.0002; \*\**p* < 0.002.

Both inoculum doses of SPONV caused rapid fatal infection. Mice in both groups were humanely euthanized starting on ~7 dpi when they appeared moribund. 100% mortality was observed for mice inoculated with 10^2^ PFU of SPONV and 93% mortality was observed for mice inoculated with 10^3^ PFU of SPONV (Fig. 1a). In comparison, 100% mortality was observed in mice inoculated with 10^2^ PFU of ZIKV-DAK. All virus inoculated groups showed significant mortality as compared to PBS-inoculated, age-matched controls (Log-rank test, *p* < 0.002) (Fig 1b). Survival curves for the group inoculated with 10^3^ PFU SPONV and the ZIKV-DAK group did not significantly differ (*p* = 0.304), whereas survival time after ZIKV-DAK infection was slightly shorter than survival time post-SPONV 10^2^ PFU infection (*p* = 0.023). Regardless of dose, SPONV-inoculated mice exhibited signs of illness by 5 to 6 dpi. We paid particular attention to the footpad used for inoculation because Salazar et al. (2019) recently noted ipsilateral footpad swelling in mice treated with an IFNAR1-blocking mAb and SPONV inoculation ^21^. However, we did not observe any notable swelling of the ipsilateral footpad during the course of our studies. The typical signs of illness we observed were more consistent with ZIKV-associated clinical signs, including weight loss, lethargy, hunched posture, and unilateral hindlimb paralysis ^29^.

### SPONV causes fetal demise in a vertical transmission mouse model

To characterize the range of pathogenic outcomes of congenital SPONV infection, we used a previously established murine pregnancy model for ZIKV ^26–28^, in which *Ifnar1*^−/−^ dams were crossed with wildtype sires to produce heterozygous offspring with one intact *Ifnar1* allele to more closely mimic the immune status of a human fetus, i.e., the fetus and placenta have intact IFN-ɑ/β signaling. Timed-mated dams were s.c. inoculated in the footpad on embryonic day 7.5 (E7.5) with 10^2^ PFU of SPONV or 10^2^ PFU ZIKV-DAK. Based on our preliminary experiments with SPONV in nonpregnant animals, and the results from our past studies ^26^, we chose this dose to minimize the potential confounding impacts of maternal illness on fetal outcomes. We collected serum samples from dams at 2 and 4 dpi to confirm maternal infection. All dams were productively infected, with detectable viremia for all groups by 4 dpi (Fig. 2a). ZIKV-DAK replicated to significantly higher titers at 4 dpi as compared to SPONV (Student’s t-test *p*-- value = 0.0008, *t* = 5.641, *df* = 7). Dams were monitored daily for clinical signs until the time of necropsy at E14.5, 7 days after inoculation. Mild clinical signs were evident in both SPONV- and ZIKV-infected dams at time of necropsy and included reduced activity, squinted eyes, ruffled fur, and hunched posture. Interestingly, clinical signs were milder in comparison to nonpregnant animals that received an equivalent inoculum dose, and none of the pregnant animals met euthanasia criteria.

**Figure 2.**
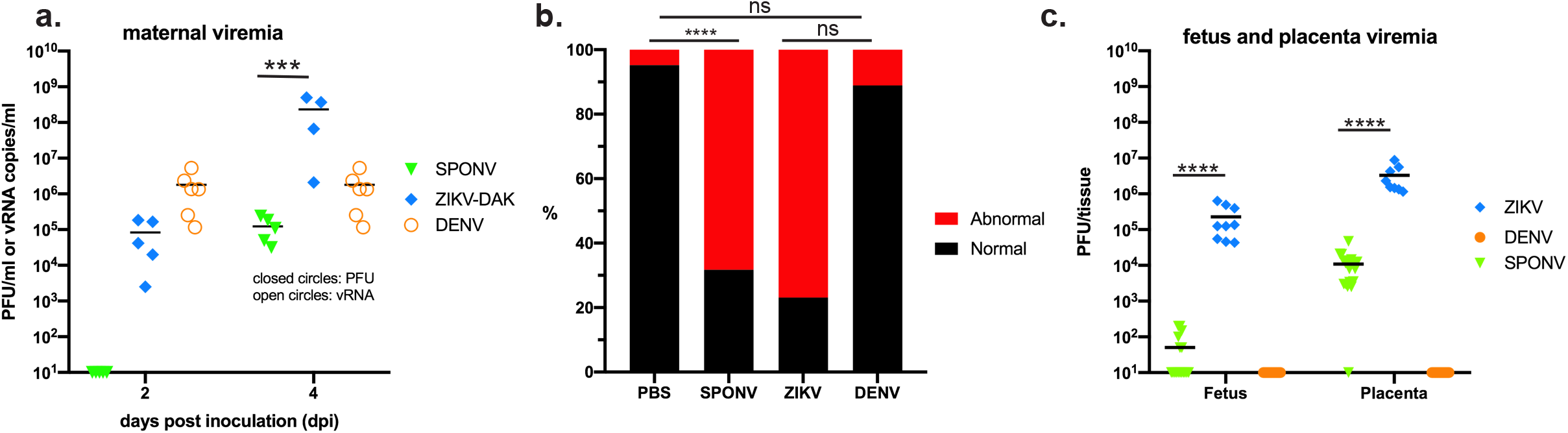
Outcomes after maternal infection at E7.5. **(a)** Time-mated *Ifnar1*^−/−^ dams were inoculated with 10^2^ SPONV or ZIKV-DAK or 10^3^ PFU of DENV at E7.5. Maternal infection with SPONV and ZIKV was confirmed by plaque assay on days 2 and 4 post inoculation (closed circles). Maternal infection with DENV was confirmed by QRT-PCR (open circles). The dotted line indicates the plaque assay limit of detection. ZIKV-DAK infected dams had significantly higher viremia at 4dpi than SPONV infected dams. \*\*\**p*< 0.002 (Student’s t-test).*­* - **(b)** Rate of grossly normal (black) versus abnormal (red) fetuses at E14.5 after maternal infection at E7.5. An abnormal fetus was defined as in any stage of resorption. Data presented are for individual fetuses from 4-6 litters per treatment group. \*\*\*\**p* < 0.0001; ns, not significant (Fisher’s exact test). **(c)** Viral titer was measured by plaque assay for a subset of individual homogenized fetuses and placentas. Symbols represent individual fetuses and placentas from 5-6 individual experiments (litters) from each treatment group. Bars represent the mean viral burden from each treatment group. No virus was detected in samples from the DENV group. Viral titers were significantly higher ZIKV placentas and fetuses as compared to SPONV. \*\*\*\**p* < 0.0001 (one-way ANOVA).

Next, to assess fetal outcomes, dams were necropsied on E14.5. Similar to what we have reported previously with different strains of ZIKV ^26^, gross examination of fetuses and placentas at time of necropsy revealed overt differences among fetuses within pregnancies and with uninfected counterparts. In general, fetuses appeared either grossly normal or abnormal, defined as being prone to embryo resorption (Fig. 2b) ^30^. At the time of necropsy, we observed high rates of resorption from both ZIKV-DAK- and SPONV-infected pregnancies. Resorption rates from ZIKV-DAK- and SPONV-infected pregnancies were not significantly different (ZIKV-DAK: 76.92% vs. SPONV: 68.29%, Fisher’s exact test, *p* = 0.457). Resorption rates for both SPONV and ZIKV-DAK were significantly higher than PBS-inoculated controls (*p* <0.0001). Despite significantly higher maternal viremia observed at 4 dpi with ZIKV-DAK-infected dams, the fact that resorption rates did not significantly differ between the two groups indicates that both ZIKV-DAK and SPONV have a propensity to harm the developing fetus that is independent of the amount of replication in maternal blood.

### Fetal harm is specific to infection with ZIKV and SPONV

Surprisingly, and in contrast to the results described by Salazar et al. where no fetal demise was observed, maternal SPONV infection in our model resulted in high rates of fetal demise ^21^. It is possible that the differences in outcomes in our study relative to Salazar et al. may be due to the use of mAb treatment versus knockout lines to abrogate interferon responses. Still, out of concern that the phenotype observed in our model was instigated by IFN-ɑ/β signaling at the maternal-interface ^27,31^ that was not ZIKV- or SPONV-specific, we inoculated *Ifnar1*^−/−^ pregnant mice with dengue virus serotype 2 (DENV-2) on E7.5. DENV is a flavivirus that is closely related to both ZIKV and SPONV and it is not known to cause adverse pregnancy outcomes in humans. To examine whether maternal DENV-2 infection is sufficient to induce fetal resorption, we s.c. inoculated pregnant dams on E7.5 with 10^3^ PFU of DENV-2. Prior to studies in pregnant animals we confirmed that this route and dose would result in productive infection in nonpregnant animals (Supplemental Figure 1). All dams were productively infected with DENV-2 with detectable vRNA loads at 2 and 4 dpi (Fig. 2a). Importantly, fetuses continued to develop as examined on E14.5, and rates of resorption were not significantly different when compared to PBS-inoculated controls (DENV-2: 11.1%, PBS: 4.8%, Fisher’s exact test, *p ­*= 0.665) (Fig. 2b). These observations confirm that fetal harm was specifically associated with ZIKV and SPONV infection. The DENV-2 results are consistent with recent findings showing that a polyinosinic/polycytidylic acid (poly[I:C], a dsRNA viral mimic) challenge could induce fetal resorption in wildtype dams but not in *Ifnar1*^−/−^ dams ^27^. These results demonstrated that peripheral maternal interferon responses could induce fetal resorption in response to nonspecific virus infection, but in *Ifnar1*^−/−^ dams, IFN-ɑ/β signalling at the maternal-fetal interface contributed to ZIKV pathogenesis.

### SPONV is vertically transmitted and causes placental and fetal histopathology

To confirm vertical transmission of SPONV a subset of placentas and fetuses were collected for plaque assay at time of necropsy from all three virus treatment groups. Infectious virus was detected in 100% of ZIKV-DAK placentas and fetuses screened (Fig. 2c). Virus was detected in all but one SPONV placenta and 35% of fetuses (Fig. 2c). No infectious virus was detected in any fetal or placental tissues following maternal DENV-2 infection at E7.5. Viral titers were significantly higher in SPONV placentas than their corresponding fetuses (one-way ANOVA with Tukey’s multiple comparisons; *p* < 0.0001), as were ZIKV-DAK placenta viral titers as compared to ZIKV-DAK fetuses (*p* = 0.006). In addition, ZIKV placental and fetal viral titers were significantly higher than SPONV titers (*p* < 0.0001)

To better understand the impact of *in utero* SPONV exposure, tissues from the developing placenta, decidua, and fetus were evaluated microscopically. In PBS- and DENV-inoculated dams, we observed normal decidua, junctional zone, and labyrinth with normal maternal and fetal blood spaces (Fig. 3). In contrast, ZIKV- and SPONV-inoculated dams displayed varying degrees of placental pathology with severe effects predominantly observed in the the labyrinth zone, including vascular injury involving maternal and/or fetal vascular spaces, infarction (obstructed blood flow), necrosis, apoptosis, and hemorrhage (Fig. 3). Overall, the severity of the vascular injury in the labyrinth zone was similar between ZIKV-DAK and SPONV placentas.

**Figure 3.**
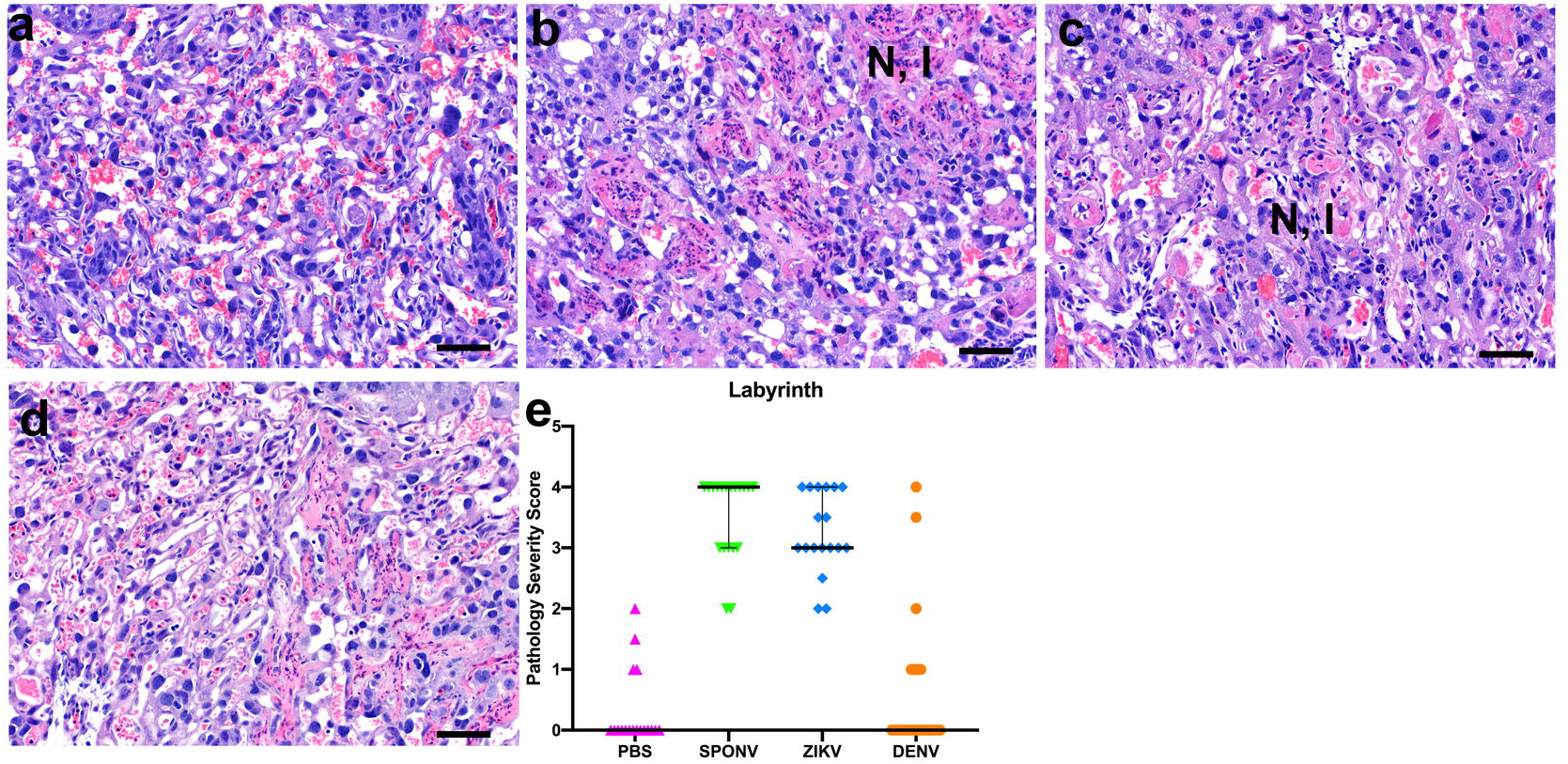
Placenta histopathology analysis: Hematoxylin and eosin (H&E) staining of placenta labyrinth zone. **(a)** Normal histologic features of the labyrinth zone of the placenta at E14.5 from a PBS inoculated dam. Necrosis (N) and inflammation (I) was observed in the labyrinth zone from SPONV **(b)** and ZIKV **(c)** inoculated dams. DENV **(d)** showed no significant placental pathology in the labyrinth zone as compared to PBS inoculated controls. Scale bar **(a-d)**, 50 μm. **(e)** The degree of placental pathology was rated on a scale of 0–4 based on the overall percent of vascular injury and/or loss in both the fetal and maternal vascular spaces of the labyrinth zone: 0 represents 0-5% vasculature injured and/or lost and 4 represents >50% injury/loss. Error bars represent 95% confidence interval from the median. Data are representative of 3–6 independent experiments for each treatment group.

In the fetuses, there was no significant microscopic pathology from PBS- and DENV-inoculated dams. In contrast, fetuses from ZIKV- and SPONV-inoculated dams demonstrated varying degrees of pathology. In fetuses from the SPONV-inoculated dams, fetal injury was evident as mild pulmonary inflammation and mild to moderate, segmental necrosis of the brain and spinal cord (Fig. 4). Pathologic findings were more widespread and severe in fetuses from ZIKV-inoculated dams and included severe necrosis and inflammation of the lung, liver, kidney, brain, and spinal cord.

**Figure 4.**
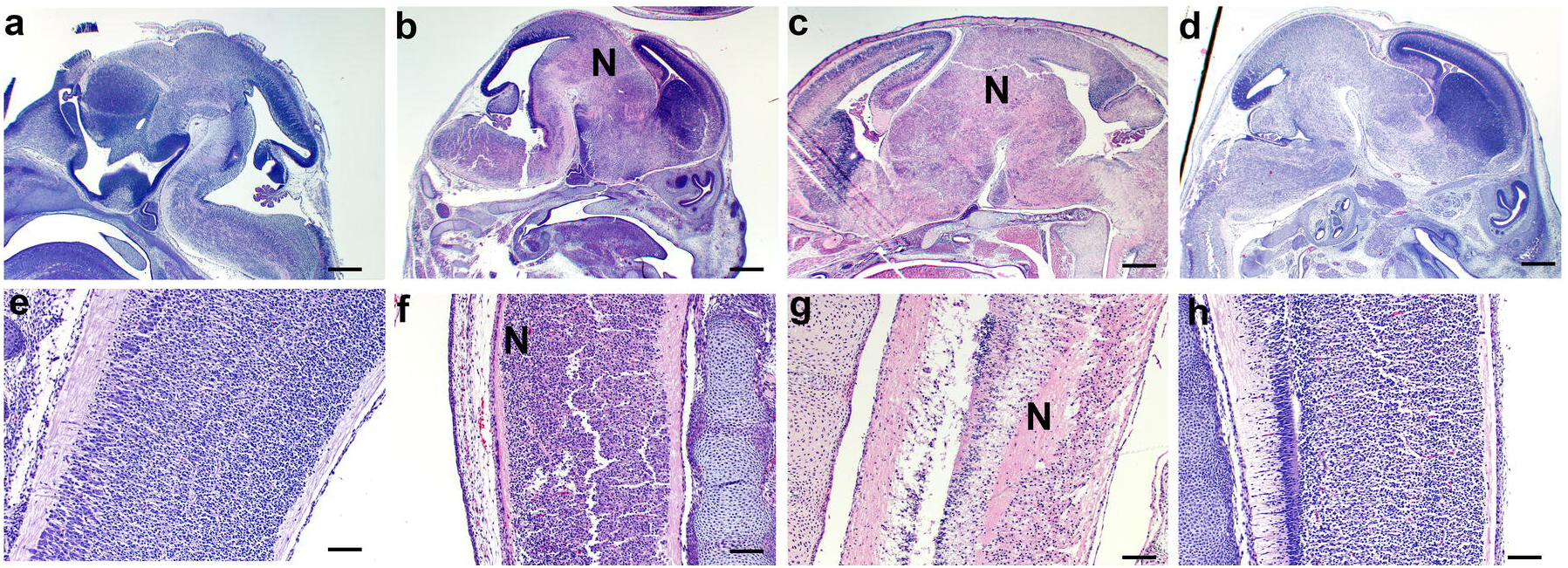
Fetus histopathology analysis: Hematoxylin and eosin (H&E) staining of fetal brain and spinal cord. **(a-d)** Fetal brains at E14.5 from PBS **(a)**, SPONV **(b)**, ZIKV **(c)**, and DENV **(d)** inoculated dams. PBS and DENV brains showed no signs of pathology. Significant histopathologic injury was observed in the fetal brains from SPONV and ZIKV inoculated dams, as indicated by large necrotic regions (N) in the brain. Scale bar **(a-d)**, 500μm. **(e-h)** Spinal cord at E14.5 from PBS **(e)**, SPONV **(f)**, ZIKV **(g)**, and DENV **(h)** inoculated dams. Fetal spinal cords from ZIKV and SPONV infections exhibited histopathologic injury as indicated by necrosis (N). Scale bar **(e-h)**, 100μm.

### *Aedes aegypti* transmits SPONV whereas *Culex quinquefasciatus* does not

To understand the potential risk for SPONV spillover into an urban epidemic cycle and because SPONV RNA was detected in a pool of *Cx. quinquefasciatus* in Haiti, we compared the relative abilities of *Ae. aegypti* and *Cx. quinquefasciatus* from Florida to transmit SPONV in the laboratory. SPONV titers in naturally infected hosts—to which feeding mosquitoes might be exposed in nature—are undefined. Therefore, we conducted our experiments with blood meal titers ranging from ~10^6^-10^8^ PFU/ml. We considered these doses to be physiologically relevant based on studies with DENV (50% mosquito infectious doses = 10^5.68^-10^7.21^ viral cDNA copies/ml) ^32^ and ZIKV (50% mosquito infectious doses = 10^6.1^-10^7.5^ PFU/ml) ^33^. To assess vector competence, mosquitoes were exposed to viremic bloodmeals via water-jacketed membrane feeder maintained at 36.5°C. Infection, dissemination, and transmission rates were assessed at 7 and 14 days post feeding (dpf) using an *in vitro* transmission assay ^34–36^. *Ae. aegypti* were susceptible to infection (10-39%) and had moderate transmission rates with all three bloodmeal concentrations (8-36%) at both 7 and 14 dpf (Table 1). Infection efficiency indicates the proportion of mosquitoes with virus-positive bodies among the tested ones. Dissemination efficiency indicates the proportion of mosquitoes with virus-positive legs, and transmission efficiency indicates the proportion of mosquitoes with infectious saliva among the tested ones. In contrast, *Cx. quinquefasciatus* were susceptible to infection (0-6%) and some of these infected mosquitoes had disseminated infections (2-3%). However, none of the infected *Cx. quinquefasciatus* were capable of transmitting virus by 14 dpf (Table 1). *Ae. aegypti* that had been exposed to ZIKV-DAK at the same time demonstrated that the assay was capable of detecting virus-positive mosquitoes at these bloodmeal concentrations.

**Table 1.** SPONV vector competence of *Aedes aegypti* and *Culex quinquefasciatus* mosquitoes. I, % infected; D, % disseminated; T, % transmitting

| Mosquito species | Virus | Dose, log<sub>10</sub> (PFU/ml) | 7 days post exposure |  |  | 14 days post exposure |  |  |
| --- | --- | --- | --- | --- | --- | --- | --- | --- |
|  |  |  | I | D | T | I | D | T |
| *Ae. aegypti* | ZIKV-DAK | **8.4-8.5 | 44/78<br>(56) | 23/40<br>(58) | 4/40<br>(10) | 33/40<br>(84) | 33/40<br>(84) | 31/40<br>(78) |
| *Ae. aegypti* | SPONV | 8.2 | 26/75<br>(35) | 23/75<br>(31) | 9/75<br>(12) | 27/70<br>(39) | 25/70<br>(36) | 25/70<br>(36) |
| *Ae. aegypti* | SPONV | 7.3 | 13/40<br>(33) | 10/40<br>(25) | 1/40<br>(3) | 13/40<br>(33) | 13/40<br>(33) | 7/40<br>(18) |
| *Ae. aegypti* | SPONV | 5.9 | 6/40<br>(15) | 3/40<br>(8) | 1/40<br>(3) | 4/40<br>(10) | 4/40<br>(10) | 3/40<br>(8) |
| *Cx. quinquefasciatus* | SPONV | 7.8 | 0/40<br>(0) | ---- | ---- | 2/40<br>(5) | 1/40<br>(3) | 0/40<br>(0) |
| *Cx. quinquefasciatus* | SPONV | 7.2 | 2/28<br>(7) | 1/28<br>(3) | 0/28<br>(0) | 1/16<br>(6) | 1/16<br>(6) | 0/16<br>(0) |
| *Cx. quinquefasciatus* | SPONV | 7.3 | 2/40<br>(5) | 1/40<br>(3) | 0/40<br>(0) | 0/40 | --- | --- |

\*I, % Infection; D, % Disseminated (of total exposed); T, % Transmitting (of total exposed).
\*\* Ranges indicate two independent experiments.

## Discussion

SPONV is the flavivirus most closely related to ZIKV and it appears to be spreading beyond Africa ^5^, but little is known about its biology. The potential of SPONV to emerge and cause congenital infections in humans as ZIKV did depends on its capacity for vertical transmission in mammals and its ability to be transmitted by human-preferring mosquitoes like *Ae. aegypti*. To date, only limited attempts have been made to evaluate its pathogenic potential and identify potential vector mosquitoes, e.g., ^21,24,25^. Here, we characterized congenital SPONV infection in a mouse model and evaluated vector competence for SPONV in two mosquito species—*Ae. aegypti* and *Cx. quinquefasciatus*—that feed on humans and are ubiquitous throughout the tropics. *Cx. quinquefasciatus* was evaluated primarily because SPONV RNA was detected in a pool of these mosquitoes during routine arbovirus surveillance activities in Haiti in 2016 ^5^. Our results demonstrated that maternal infection with SPONV caused significant fetal harm consistent with ZIKV infections in this same model. Importantly, we confirmed that outcomes were specific to SPONV and ZIKV infection, because we observed no fetal harm in dams infected with DENV-2. Although there are limitations regarding the translational relevance of this model, this same mouse platform is widely regarded as establishing the first direct causal link between ZIKV and fetal abnormalities in 2016 ^28^. We therefore speculate that SPONV infection during pregnancy in humans poses a risk to the developing neonate, and suggest that clinicians in areas endemic for *Ae. aegypti* caring for non-ZIKV adverse outcomes consider the possibility of congenital SPONV infection. We also speculate that SPONV could follow in the path of other *Ae. aegypti*-transmitted arboviruses, because of our data demonstrating that *Ae. aegypti*—which also transmits DENV, ZIKV, yellow fever virus and chikungunya virus—could transmit SPONV. *Ae. aegypti* is a highly domesticated urban mosquito that lives in close proximity to humans, feeds on humans, and lays eggs in containers made by humans ^37^. As a result, *Ae. aegypti* thrives in conditions created by overcrowding in tropical cities and thus facilitates epidemic arbovirus transmission.

In this study we found that SPONV infection of nonpregnant *Ifnar1*^−/−^ mice was nearly uniformly lethal, which mirrors observations from other studies using AG129 mice, which lack both type I and II interferon receptors ^24^, and mice treated with an anti-IFNAR1 mAb to transiently abrogate type I interferon signaling ^21^. However, mice succumbed significantly faster in our model compared to mAb-treated mice. This is consistent with observations of ZIKV infection, which also results in less mortality in the mAb model as compared to interferon-deficient animals ^29^. Surprisingly, pregnant animals that received the anti-IFNAR1 mAb treatment prior to SPONV inoculation showed infection of the placenta and vertical transmission but not increased rates of fetal resorption ^21^. We also found evidence of placental infection and vertical transmission. In contrast, SPONV-inoculated dams had significant rates of fetal resorption and severe pathology at both the maternal-fetal interface and in the developing fetus in our study, consistent with ZIKV-inoculated controls and with what we and others have reported previously for ZIKV ^26–28^.

It is possible that differences in fetal outcomes and histopathology observed in our study relative to the Salazar et al. study may be due to the use of a different strain, dose, or mouse platform. It should be noted, however, that our *Ifnar1*^−/−^ mice are on the same genetic background as the mice used in the Salazar study (C57BL/6), therefore differences in outcomes are not likely to be due to variation in host loci outside of the IFNAR1 gene. In addition, Salazar et al. used the same SA Ar94 strain used in our study (SRA accession number pending), but virus stocks prepared by different laboratories, while nominally the same strain, have different passage histories, which could result in small, but biologically important, phenotypic differences, e.g., ^38^. Another possible explanation for the disparity in outcomes is the mouse platform. The *Ifnar1^+/-^* platform we used has consistently resulted in more severe outcomes in previous studies with ZIKV, whereas the anti-IFNAR1 mAb platform has demonstrated ZIKV replication and detection in the placenta and fetus-the extent of which was dependent on dose of mAb used ^28^. Importantly, in the Minner et al. study demise was not observed when anti-IFNAR1 mAb was administered prior to ZIKV inoculation but it was observed when *Ifnar1*^−/−^ dams were used ^28^. These data appear to be consistent with SPONV outcomes using the same platforms (^21^ and results described herein).

Despite similar fetal outcomes, SPONV and ZIKV may not necessarily utilize the same pathways or molecular mechanisms to harm the fetus. Histopathological analyses demonstrated significant fetal and placental pathology in SPONV- and ZIKV-inoculated groups as compared to PBS- or DENV-inoculated controls. But ZIKV-DAK-inoculated dams displayed the most severe histologic phenotype that corresponded with higher placenta and fetus titers (Fig 2). SPONV histopathology was more heterogeneous and severity scores were reminiscent of what we have previously observed with congenital Asian-lineage ZIKV infection ^26^. The labyrinth zones of both ZIKV and SPONV placentas showed similar degrees of vascular injury/loss, such that, despite this heterogeneity, the final outcome of fetal demise did not differ between these two viruses. Importantly, we only assessed outcomes following a single infection timepoint at E7.5 and infection outcomes may differ depending on the timing of infection ^39^. More studies are needed to better understand antiviral signalling at the maternal fetal interface and pathways these viruses exploit to traffic to the feto-placental unit to cause harm during pregnancies. Translational nonhuman primate models will be valuable for such studies.

SPONV’s ability to cause wide-scale human infections depends on the mosquito vector that transmits it. One recent study suggested that strains of *Ae. aegypti*, *Ae. albopictus*, and *Cx. quinquefasciatus* were refractory to SPONV infection ^25^. However, the blood on which mosquitoes fed in these experiments harbored moderate SPONV infectivity of ~10^5^ PFU/ml. For related arboviruses like DENV, it is known that the level of viremia is one of the most important determinants of human infectiousness to mosquitoes ^40,41^. ZIKV 50% mosquito infectious doses are in the 10^6.1^-10^7.5^ PFU/ml range ^33^. If SPONV is similar, then studies employing bloodmeal titers of ~10^5^ PFU/ml are too low. In addition, some studies have shown that freeze-thawing flaviviruses impacts mosquito infection and transmission rates ^3503398,28430564^. As a result, we chose a virus dose that we estimated to be what a mosquito might encounter in nature based on previous studies with DENV and ZIKV ^32,33^, the flaviviruses most closely related to SPONV. In addition, we chose to prepare our infectious bloodmeals using fresh virus supernatant to avoid the confounding effects of freeze-thawing virus stocks. These experiments demonstrated that *Ae. aegypti* could be productively infected and could transmit SPONV-as indicated by infectious virus in the saliva, suggesting the ability to transmit to a vertebrate host. In contrast, *Cx. quinquefasciatus* were susceptible to infection but none of these infected mosquitoes were capable of transmitting virus by day 14 post-feeding. These data therefore suggest that the wild-caught Haiti mosquitoes likely contained trace amounts of viremic blood in their midguts indicating that they fed upon a SPONV-infected human or animal. Further, the Haiti data also are consistent with field epidemiological reports for which it has been difficult to detect DENV-, CHIKV-, and/or ZIKV-infected mosquitoes because of low infection rates in field-collected mosquitoes, even during periods of active human outbreaks ^43–45^. Finally, we used only a single combination of mosquito and virus genotype. Vector competence of many mosquito species for certain viruses likely is governed by vector genotype x virus genotype interactions in genetically diverse, natural mosquito populations (reviewed in ^46^). These data at the very least argue for more studies—both experimental and epidemiological—assessing different SPONV-mosquito combinations.

Together, our data show that SPONV posessess worrisome properties that should prompt further investigation into its epidemic potential and risk to pregnant women. The adaptation of SPONV to an urban or peri-urban cycle, involving *Ae. aegypti* and/or other mosquitoes in the *Stegomyia* subgenus (e.g., *Ae. albopictus*) should therefore be a public health concern ^47^. Given the continuing difficulties in differentiating between flaviviruses in diagnostic assays, understanding SPONV’s prevalence in the expanding landscape of cross-reacting, co-endemic mosquito-borne viruses could also be considered a critical public health priority.

## Methods

### Ethical Approval

This study was approved by the University of Minnesota, Twin Cities Institutional Animal Care and Use Committee (Protocol Number 1804–35828).

### Cells and viruses

African Green Monkey kidney cells (Vero; ATCC #CCL-81) were maintained in Dulbecco’s modified Eagle medium (DMEM) supplemented with 10% fetal bovine serum (FBS; Hyclone, Logan, UT), 2 mM L-glutamine, 1.5 g/L sodium bicarbonate, 100 U/ml penicillin, 100 μg/ml of streptomycin, and incubated at 37°C in 5% CO2. *Aedes albopictus* mosquito cells (C6/36; ATCC #CRL-1660) were maintained in DMEM supplemented with 10% fetal bovine serum (FBS; Hyclone, Logan, UT), 2 mM L-glutamine, 1.5 g/L sodium bicarbonate, 100 U/ml penicillin, 100 μg/ml of streptomycin, and incubated at 28°C in 5% CO2. The cell lines were obtained from the American Type Culture Collection, were not further authenticated, and were not specifically tested for mycoplasma.

ZIKV strain DAK AR 41524 (ZIKV-DAK; GenBank:KY348860) was originally isolated from *Aedes africanus* mosquitoes in Senegal in 1984, with a round of amplification on *Aedes pseudocutellaris* cells, followed by amplification on C6/36 cells, followed by two rounds of amplification on Vero cells, was obtained from BEI Resources (Manassas, VA). SPONV strain SA Ar94 (GenBank: KX227370) was originally isolated from a *Mansonia uniformis* mosquito in Lake Simbu, Natal, South Africa in 1955, with five rounds of amplification with unknown culture conditions followed by a single round of amplification on Vero cells. Virus stocks were prepared by inoculation onto a confluent monolayer of C6/36 mosquito cells. Contemporary SPONV isolates from Haiti do not exist. The prototype Chuku strain (Nigeria, 1952) has an unknown passage history. We choose not to use such an isolate for *in vivo* studies - we previously showed that the prototype ZIKV strain MR766, which had undergone extensive passaging in suckling mice, was attenuated in monkeys ^48^. Instead we used the only available low-passage isolate, SPONV strain SA Ar 94. This strain is 98.8% nucleotide identical with the SPONV genome recovered from mosquitoes in Haiti (Genbank:MG182017), but we acknowledge that even though the sequences are almost identical, the slight difference could result in important phenotypic impacts. DENV-2 strain BID-V594 (GenBank:EU482725), originally isolated from a human in Puerto Rico in 2006 with a single round of amplification on C6/36 cells, was obtained from BEI Resources (Manassas, VA). BEI amplified the virus on C6/36 cells and virus stocks were prepared by inoculation onto a confluent monolayer of C6/36 mosquito cells. We deep sequenced our virus stocks to verify the expected origin (see next section for details). The SPONV, ZIKV-DAK, and DENV-2 stocks matched the GenBank sequences (MG182017, KY348860, EU482725, respectively) of the parental viruses; but a single nucleotide position in the 3’ UTR (site 10,603) of the DENV-2 stock contained a 79/21 ratio of T-to-A nucleotide substitutions and a variant at site 3710 in the ZIKV-DAK stock encodes a nonsynonymous change (A to V) in NS2A.

### Deep Sequencing

A vial of the viral stocks used for primary challenge (ZIKV-DAK, SPONV, DENV-2), were each deep sequenced by preparing libraries of fragmented double-stranded cDNA using methods similar to those previously described ^49^. Briefly, the sample was centrifuged at 5000 rcf for five minutes. The supernatant was then filtered through a 0.45-μm filter. Viral RNA was isolated using the QIAamp MinElute Virus Spin Kit (Qiagen, Germantown, MD), omitting carrier RNA. Eluted vRNA was then treated with DNAse I. Double-stranded DNA was prepared with the Superscript Double-Stranded cDNA Synthesis kit (Invitrogen, Carlsbad, CA) and priming with random hexamers. Agencourt Ampure XP beads (Beckman Coulter, Indianapolis, IN) were used to purify double-stranded DNA. The purified DNA was fragmented with the Nextera XT kit (Illumina, Madison, WI), tagged with Illumina-compatible primers, and then purified with Agencourt Ampure XP beads. Purified libraries were then sequenced with 2 x 300 bp kits on an Illumina MiSeq.

### Sequence Analysis

Viral stock sequences were analyzed using a modified version of the viral-ngs workflow developed by the Broad Institute (http://viral-ngs.readthedocs.io/en/latest/description.html) implemented in DNANexus and using bbmap local alignment in Geneious Pro (Biomatters, Ltd., Auckland, New Zealand). Briefly, using the viral-ngs workflow, host-derived reads that map to a human sequence database and putative PCR duplicates were removed. The remaining reads were loaded into Geneious Pro and mapped to NCBI Genbank Zika (GenBank:KX601166), Spondweni (Genbank:MG182017), or dengue virus (GenBank:EU482725) reference sequences using bbmap local alignment. Mapped reads were aligned using Geneious global alignment and the consensus sequence was used for intra sample variant calling. Variants were called that fit the following conditions: have a minimum *p*-value of 10e-60, a minimum strand bias of 10e-5 when exceeding 65% bias, and were nonsynonymous.

### Plaque assay

All ZIKV and SPONV screens from mouse tissue and titrations for virus quantification from virus stocks were completed by plaque assay on Vero cell cultures. Duplicate wells were infected with 0.1ml aliquots from serial 10-fold dilutions in growth media and virus was adsorbed for one hour. Following incubation, the inoculum was removed, and monolayers were overlaid with 3ml containing a 1:1 mixture of 1.2% oxoid agar and 2X DMEM (Gibco, Carlsbad, CA) with 10% (vol/vol) FBS and 2% (vol/vol) penicillin/streptomycin. Cells were incubated at 37°C in 5% CO2 for four days for plaque development for ZIKV and five days for SPONV. Cell monolayers then were stained with 3ml of overlay containing a 1:1 mixture of 1.2% oxoid agar and 2X DMEM with 2% (vol/vol) FBS, 2% (vol/vol) penicillin/streptomycin, and 0.33% neutral red (Gibco). Cells were incubated overnight at 37°C and plaques were counted.

### Viral RNA isolation

DENV-2 viral RNA was extracted from sera using the Viral Total Nucleic Acid Kit (Promega, Madison, WI) on a Maxwell 48 RSC instrument (Promega, Madison, WI). Viral RNA was isolated from homogenized tissues using the Maxwell 48 RSC Viral Total Nucleic Acid Purification Kit (Promega, Madison, WI) on a Maxwell 48 RSC instrument. Each tissue was homogenized using PBS supplemented with 20% FBS and penicillin/streptomycin and a tissue tearor variable speed homogenizer. Supernatant was clarified by centrifugation and the isolation was continued according to the Maxwell 48 RSC Viral Total Nucleic Acid Purification Kit protocol, and samples were eluted into 50 μl RNase free water. RNA was then quantified using quantitative RT-PCR. Viral load data from serum are expressed as vRNA copies/mL. Viral load data from tissues are expressed as vRNA copies/tissue.

### Quantitative reverse transcription PCR (QRT-PCR)

For DENV-2, vRNA from serum and tissues was quantified by QRT-PCR using the following primers: Forward: CAGATCTCTGATGAAYAACCAACG Reverse: AGTYGACACGCGGTTTCTCT Probe: 6-Fam-CGCGTTTCAGCATATTGAA-BHQ1. IUPAC nucleotide codes are as follows: Y: C or T; B: C or G or T; R: A or G. The RT-PCR was performed using the SuperScript III Platinum One-Step Quantitative RT-PCR system (Invitrogen, Carlsbad, CA) on a LightCycler 480 instrument (Roche Diagnostics, Indianapolis, IN).The primers and probe were used at final concentrations of 600 nm and 100 nm respectively, along with 150 ng random primers (Promega, Madison, WI). Cycling conditions were as follows: 37°C for 15 min, 50°C for 30 min and 95°C for 2 min, followed by 50 cycles of 95°C for 15 sec and 60°C for 1 min. Viral RNA concentration was determined by interpolation onto an internal standard curve composed of seven 10-fold serial dilutions of a synthetic DENV-2 RNA fragment based the WHO type strain, New Guinea C that shares ~94% similarity at the nucleotide level to the Puerto Rican strain used in the infections described in this manuscript.

### Mice

Female *Ifnar1*^−/−^ mice on the C57BL/6 background were bred in the specific pathogen-free animal facilities of the University of Minnesota College of Veterinary Medicine. Male C57BL/6 mice were purchased from Jackson Laboratories. Timed matings between female *Ifnar1*^−/−^ mice and male C57BL/6 mice resulted in *Ifnar1^−/+^* progeny.

### Subcutaneous inoculation

Non-pregnant mice were between six and eleven weeks of age. All pregnant dams were between six and nine weeks of age. Littermates were randomly assigned to infected and control groups. Matings between female *Ifnar1*^−/−^ dams and wildtype sires were timed by checking for the presence of a vaginal plug, indicating a gestational age E0.5. At embryonic day 7.5 (E7.5), dams were inoculated in the left hind foot pad with 10^2^ PFU of ZIKV or SPONV, in 20 μl of sterile PBS, with 10^3^ PFU of DENV in 25μl of sterile PBS, or with 25 μl of sterile PBS alone to serve as experimental controls. All animals were closely monitored by laboratory staff for adverse reactions and signs of disease. Sub-mandibular blood draws were performed 2 and 4 days post inoculation and serum was collected to verify viremia. Mice were humanely euthanized and necropsied at E14.5.

### Mouse necropsy

Following inoculation with SPONV, ZIKV, or PBS, mice were sacrificed at E14.5. Tissues were carefully dissected using sterile instruments that were changed between each mouse to minimize possible cross contamination. For all mice, each organ/neonate was evaluated grossly *in situ*, removed with sterile instruments, placed in a sterile culture dish, and further processed to assess viral burden and tissue distribution or banked for future assays. Briefly, uterus was first removed, and then dissected to remove each individual conceptus (i.e, fetus and placenta when possible). Fetuses and placentas were either collected in PBS supplemented with 20% FBS and penicillin/streptomycin (for plaque assays) or fixed in 4% PFA or 10% Neutral Buffered Formalin for imaging. We characterized an embryo as in the resorption process if it met the following criteria: significant growth retardation compared to litter mates and controls accompanied by clearly evident developmental delay, i.e., morphology was ill defined; or visualization of a macroscopic plaque in the uterus ^30^.

### Histology

Tissues were fixed in 4% paraformaldehyde for 24 hours and transferred into cold, sterile DPBS until alcohol processed and embedded in paraffin. Paraffin sections (5 μm) were stained with hematoxylin and eosin (H&E). Pathologists were blinded to gross pathological findings when tissue sections were evaluated microscopically. The degree of pathology at the maternal-fetal interface was rated on a scale of 0-4: 0 – no lesions (normal); 1 – mild changes (1-2 focal lesions or 10-15% of zone involved); 2 – mild to moderate changes (3-4 focal lesions or 10-15% of zone involved); 3 – moderate to severe changes (4-6 focal lesions or 15-25% of zone involved); 4 – severe (>6 focal lesions or >25% of zone involved). The final score was dependent upon the greater of two parameters (# of lesions or % zone involved). This was an identical scoring system to what we reported previously ^26^. The final scores were determined as a consensus score of two independent pathologists. For each zone in the placenta (myometrium, decidua, junctional zone, labyrinth, and chorionic plate/membranes) a ‘General’ overall score was determined, a score for the amount of ‘Inflammation’, and a score for direct ‘Vascular Injury’. The ‘General’ score was based on an interpretation of the overall histopathologic findings in each placenta, which included features of necrosis, infarction, apoptosis, hemorrhage, thrombosis, mineralization, vascular injury, and inflammation. The ‘Inflammation’ score quantified the amount of inflammation in that layer. The ‘Vascular Injury’ score assessed vascular wall injury (fibrinoid necrosis, endothelial swelling), dilatation of the vessels or spaces, necrosis, loss of vascular lumen diameter, and intraluminal thrombi. The myometrial layer representing the uterine wall and the chorionic plate/membranes were often not present in histologic sections and therefore meaningful comparisons between strains could not be assessed. The decidual layer (maternal in origin), the junctional zone composed of fetal giant cells and spongiotrophoblast, and the labyrinth layer (the critical layer for gas and nutrient exchange between the fetal and maternal vascular systems) were scored. Since the percentage of injured/pathologic labyrinth zone is probably the best predictor of poor fetal outcome, we also independently scored the labyrinth zone based only on the percentage of fetal and maternal vascular injury/loss using the following scoring system: 0-5%-0 (background); 5-15%-1 (mild); 15-30%-2 (moderate); 30-50%-3 (moderate to severe); and >50%-4 (severe). Photomicrographs were obtained using a bright light microscope Olympus BX43 and Olympus BX46 (Olympus Inc., Center Valley, PA) with attached Olympus DP72 digital camera (Olympus Inc.) and Spot Flex 152 64 Mp camera (Spot Imaging), and captured using commercially available image-analysis software (cellSens DimensionR, Olympus Inc. and spot software 5.2).

### Mosquito strain and colony maintenance

All mosquitoes used in this study were maintained at the University of Minnesota, Twin Cities as described ^50^ in an environmental chamber at 26.5 ± 1 °C, 75% ± 5% relative humidity, and with a 12 hour light and 12 hour dark photoperiod with a 30 minute crepuscular period at the beginning of each light cycle. The *Aedes aegypti* Gainesville strain used in this study was obtained from Mike Smanski (University of Minnesota, Twin Cities, St. Paul, MN), and was originally derived from the USDA “Gainesville” strain. *Culex quinquefasciatus* used in this study were obtained from Greg Ebel (Colorado State University, Ft. Collins, CO) and were originally collected in Sebring County, FL in 1988. Three-to six-day-old female mosquitoes were used for all experiments.

### Vector competence studies

Mosquitoes were exposed to SPONV- or ZIKV-infected bloodmeals via water-jacketed membrane feeder maintained at 36.5 °C ^51^. Bloodmeals consisted of defibrinated sheep blood (HemoStat Laboratories, Inc.) and fresh virus supernatant, yielding infectious bloodmeal titers ranging from ~10^6^-10^8^ PFU/ml. Bloodmeal titer was determined after feeding. Infection, dissemination, and transmission rates were determined for individual mosquitoes and sample sizes were chosen using long established procedures ^35,36,52^. Briefly, mosquitoes were sucrose starved for 14 to 16 hours prior to bloodmeal exposure. Mosquitoes that fed to repletion were randomized, separated into cartons in groups of 40-50, and maintained on 0.3 M sucrose in a Conviron A1000 environmental chamber at 26.5 ± 1 °C, 75% ± 5% relative humidity, with a 12 hour photoperiod within the Veterinary Isolation Facility BSL3 Insectary at the University of Minnesota, Twin Cities. All samples were screened by plaque assay on Vero cells.

### Data analysis

All analysis were performed using GraphPad Prism. For survival analysis, Kaplan-Meier survival curves were analyzed by the log-rank test. Unpaired Student’s t-test was used to determine significant differences in maternal viremia and fetal and placental tissue viremia. Fisher’s exact test was used to determine differences in rates of normal vs. abnormal concepti.

## Acknowledgements

The authors acknowledge the University of Minnesota, Twin Cities BSL3 Program for facilities and Neal Heuss for support. We thank Natalie Benett for her contribution in mosquito maintenance, and the University of Minnesota, Twin Cities Comparative Pathology Shared Resource for preparation of histological sections. Funding for this project came from DHHS/PHS/NIH R21AI131454 and R01 AI132563 to M.T.A. The publication’s contents are solely the responsibility of the authors and do not necessarily represent the official views of the NCRR or NIH.

## Author Contributions

A.S.J. and M.T.A. designed experiments. A.S.J., S.L.O., D.H.O, D.M.S., T.C.F., and M.T.A. analyzed data and drafted the manuscript. A.S.J. and M.T.A. performed mouse and vector competence studies. A.M.W., R.S., and T.C.F. developed and performed viral load assays. R.M.V. and S.L.O. developed and performed the deep sequencing pipeline. D.M.S. and M.K.F. performed histological analysis.

## Competing Financial Interests

The authors declare no competing financial interests.

---

**Supplemental Figure 1.**
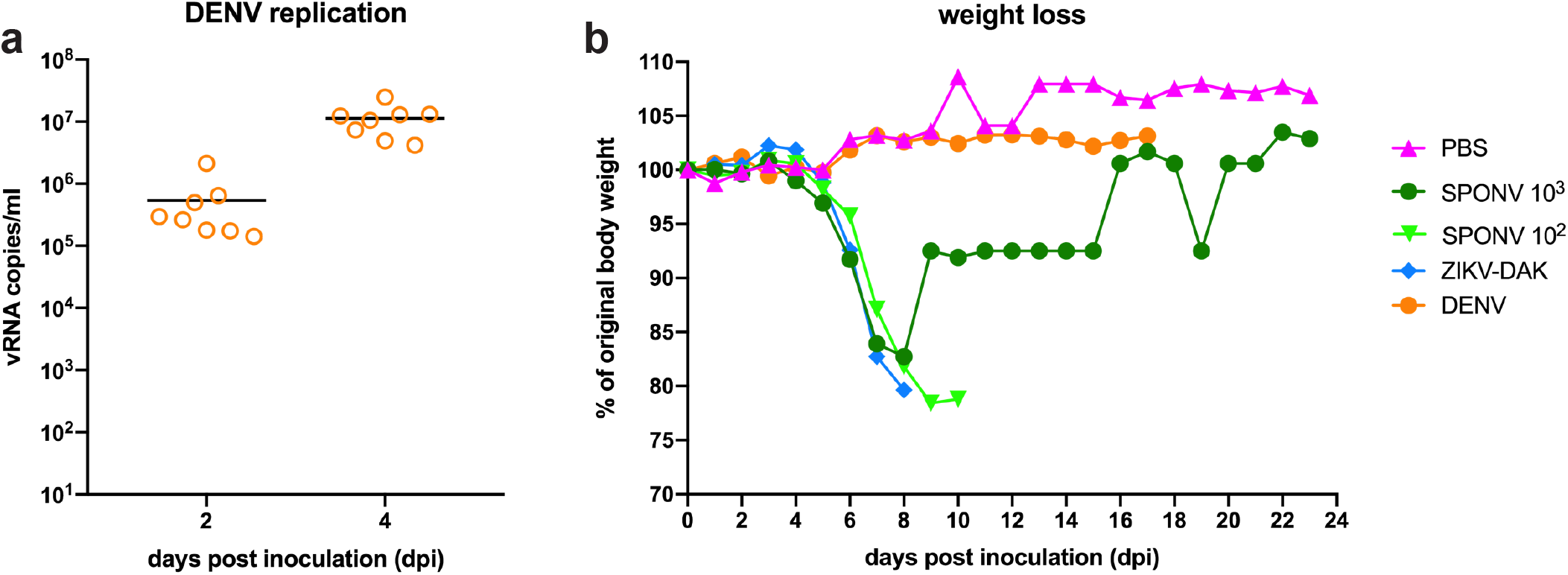
Confirmation of DENV replication and weight loss in non-pregnant *Ifnar1*^−/−^ mice. **(a)** Prior to pregnancy experiments, 8 male seven-to nine-week-old mice were s.c. inoculated with 10^3^ PFU of DENV-2 to confirm virus replication. Serum was collected 2 and 4dpi to confirm infection. Viral loads were measured by QRT-PCR. All 8 mice had detectable vRNA at both 2 and 4 dpi. No clinical signs were observed. **(b)** Six-to eleven-week-old mice were weighed daily after s.c. inoculation with PBS, SPONV, ZIKV, or DENV. The first replicate, surviving mice were monitored for 24 days post inoculation. For following replicates, mice were monitored for up to 17dpi. PBS and DENV infected mice did not lose weight or show clinical signs. A single SPONV 10^3^-inoculated mouse survived beyond 9 dpi.

## References

1. Dick, G. W., Kitchen, S. F. & Haddow, A. J. Zika virus. I. Isolations and serological specificity. Trans R Soc Trop Med Hyg 46, 509–520 (1952).

2. Simpson, D. I. Zika VIRUS INFECTION IN MAN. Trans R Soc Trop Med Hyg 58, 335–338 (1964).

3. Johansson, M. A., Mier-y-Teran-Romero, L., Reefhuis, J., Gilboa, S. M. & Hills, S. L. Zika and the Risk of Microcephaly. N Engl J Med 375, 1–4 (2016).

4. Melo, A. S. et al. Congenital Zika Virus Infection: Beyond Neonatal Microcephaly. JAMA Neurol 73, 1407–1416 (2016).

5. White, S. K., Lednicky, J. A., Okech, B. A., Morris, J. G. & Dunford, J. C. Spondweni Virus in Field-Caught Culex quinquefasciatus Mosquitoes, Haiti, 2016. Emerg Infect Dis 24, 1765–1767 (2018).

6. Wolfe, M. S., Calisher, C. H. & McGuire, K. Spondweni virus infection in a foreign resident of Upper Volta. Lancet 2, 1306–1308 (1982).

7. Theiler, M. & Downs, W. G. The arthropod-borne viruses of vertebrates. An account of theRockefeller Foundation virus program 1951–1970. 174 (1973).

8. Haddow, A. J., Williams, M. C., Woodall, J. P., Simpson, D. I. & Goma, L. K. TWELVE ISOLATIONS OF ZIKA VIRUS FROM AEDES (STEGOMYIA) AFRICANUS (THEOBALD) TAKEN IN AND ABOVE A UGANDA FOREST. Bull World Health Organ 31, 57–69 (1964).

9. Draper, C. C. INFECTION WITH THE CHUKU STRAIN OF SPONDWENI VIRUS. West Afr Med J 14, 16–19 (1965).

10. Macnamara, F. N. Zika virus: a report on three cases of human infection during an epidemic of jaundice in Nigeria. Trans R Soc Trop Med Hyg 48, 139–145 (1954).

11. Kokernot, R. H., Smithburn, K. C., Muspratt, J. & Hodgson, B. Studies on arthropod-borne viruses of Tongaland. VIII. Spondweni virus, an agent previously unknown, isolated from Taeniorhynchus (Mansonioides) uniformis. S Afr J Med Sci 22, 103–112 (1957).

12. Kokernot, R. H., Casaca, V. M., Weinbren, M. P. & McIntosh, B. M. Survey for antibodies against arthropod-borne viruses in the sera of indigenous residents of Angola. Trans R Soc Trop Med Hyg 59, 563–570 (1965).

13. Kokernot, R. H., Szlamp, E. L., Levitt, J. & McIntosh, B. M. Survey for antibodies against arthropod-borne viruses in the sera of indigenous residents of the Caprivi Strip and Bechuanaland Protectorate. Trans R Soc Trop Med Hyg 59, 553–562 (1965).

14. Brottes, H., Rickenbach, A., Brès, P., Salaün, J. J. & Ferrara, L. [Arboviruses in the Cameroon. Isolation from mosquitoes]. Bull World Health Organ 35, 811–825 (1966).

15. Ardoin, P., Rodhain, F. & Hannoun, C. Epidemiologic study of arboviruses in the Arba-Minch district of Ethiopia. Trop Geogr Med 28, 309–315 (1976).

16. Haddow, A. D. & Woodall, J. P. Distinguishing between Zika and Spondweni viruses. Bull World Health Organ 94, 711–711A (2016).

17. Mcintosh, B. M., Kokernot, R. H., Paterson, H. E. & De Meillon, B. Isolation of Spondweni virus from four species of culicine mosquitoes and a report of two laboratory infections with the virus. S Afr Med J 35, 647–650 (1961).

18. Boorman, J. P. & Draper, C. C. Isolations of arboviruses in the Lagos area of Nigeria, and a survey of antibodies to them in man and animals. Trans R Soc Trop Med Hyg 62, 269–277 (1968).

19. Platt, D. J. et al. Zika virus-related neurotropic flaviviruses infect human placental explants and cause fetal demise in mice. Sci Transl Med 10, (2018).

20. Andersen, A. A. & Hanson, R. P. Experimental transplacental transmission of st. Louis encephalitis virus in mice. Infect Immun 2, 320–325 (1970).

21. Salazar, V. et al. Dengue and Zika Virus Cross-Reactive Human Monoclonal Antibodies Protect against Spondweni Virus Infection and Pathogenesis in Mice. Cell Rep 26, 1585–1597.e4 (2019).

22. McIntosh, B. M., Jupp, P. G. & De Sousa, J. Further isolations of the arboviruses from mosquitoes collected in Tongaland, South Africa, 1960-1968. J Med Entomol 9, 155–159 (1972).

23. Worth, C. B., Paterson, H. E. & De Meillon, B. The incidence of arthropod-borne viruses in a population of culicine mosquitoes in Tongaland, Union of South Africa (January, 1956, through April, 1960). Am J Trop Med Hyg 10, 583–592 (1961).

24. McDonald, E. M., Duggal, N. K. & Brault, A. C. Pathogenesis and sexual transmission of Spondweni and Zika viruses. PLoS Negl Trop Dis 11, e0005990 (2017).

25. Haddow, A. D. et al. Genetic Characterization of Spondweni and Zika Viruses and Susceptibility of Geographically Distinct Strains of Aedes aegypti, Aedes albopictus and Culex quinquefasciatus (Diptera: Culicidae) to Spondweni Virus. PLoS Negl Trop Dis 10, e0005083 (2016).

26. Jaeger, A. S. et al. Zika viruses of African and Asian lineages cause fetal harm in a mouse model of vertical transmission. PLoS Negl Trop Dis 13, e0007343 (2019).

27. Yockey, L. J. et al. Type I interferons instigate fetal demise after Zika virus infection. Sci Immunol 3, (2018).

28. Miner, J. J. et al. Zika Virus Infection during Pregnancy in Mice Causes Placental Damage and Fetal Demise. Cell 165, 1081–1091 (2016).

29. Lazear, H. M. et al. A Mouse Model of Zika Virus Pathogenesis. Cell Host Microbe 19, 720–730 (2016).

30. Flores, L. E., Hildebrandt, T. B., Kühl, A. A. & Drews, B. Early detection and staging of spontaneous embryo resorption by ultrasound biomicroscopy in murine pregnancy. Reprod Biol Endocrinol 12, 38 (2014).

31. Casazza, R. L. & Lazear, H. M. Antiviral immunity backfires: Pathogenic effects of type I interferon signaling in fetal development. Sci Immunol 3, (2018).

32. Duong, V. et al. Asymptomatic humans transmit dengue virus to mosquitoes. Proc Natl Acad Sci U S A 112, 14688–14693 (2015).

33. Ciota, A. T. et al. Effects of Zika Virus Strain and Aedes Mosquito Species on Vector Competence. Emerg Infect Dis 23, 1110–1117 (2017).

34. Aliota, M. T., Peinado, S. A., Velez, I. D. & Osorio, J. E. The wMel strain of Wolbachia Reduces Transmission of Zika virus by Aedes aegypti. Sci Rep 6, 28792 (2016).

35. Dudley, D. M. et al. Infection via mosquito bite alters Zika virus tissue tropism and replication kinetics in rhesus macaques. Nat Commun 8, 2096 (2017).

36. Aliota, M. T., Peinado, S. A., Osorio, J. E. & Bartholomay, L. C. Culex pipiens and Aedes triseriatus Mosquito Susceptibility to Zika Virus. Emerg Infect Dis 22, 1857–1859 (2016).

37. Gubler, D. J. Dengue, Urbanization and Globalization: The Unholy Trinity of the 21(st) Century. Trop Med Health 39, 3–11 (2011).

38. Duggal, N. K. et al. Mutations present in a low-passage Zika virus isolate result in attenuated pathogenesis in mice. Virology 530, 19–26 (2019).

39. Jagger, B. W. et al. Gestational Stage and IFN-λ Signaling Regulate ZIKV Infection In Utero. Cell Host Microbe 22, 366–376.e3 (2017).

40. Nguyet, M. N. et al. Host and viral features of human dengue cases shape the population of infected and infectious Aedes aegypti mosquitoes. Proc Natl Acad Sci U S A 110, 9072–9077 (2013).

41. Carrington, L. B. & Simmons, C. P. Human to mosquito transmission of dengue viruses. Front Immunol 5, 290 (2014).

42. Miller, B. R. Increased yellow fever virus infection and dissemination rates in Aedes aegypti mosquitoes orally exposed to freshly grown virus. Trans R Soc Trop Med Hyg 81, 1011–1012 (1987).

43. Grubaugh, N. D. et al. Genomic epidemiology reveals multiple introductions of Zika virus into the United States. Nature 546, 401–405 (2017).

44. Dzul-Manzanilla, F. et al. Evidence of vertical transmission and co-circulation of chikungunya and dengue viruses in field populations of Aedes aegypti (L.) from Guerrero, Mexico. Trans R Soc Trop Med Hyg 110, 141–144 (2016).

45. Ramesh, A. et al. No evidence of Zika, dengue, or chikungunya virus infection in field-caught mosquitoes from the Recife Metropolitan Region, Brazil, 2015. Wellcome Open Res 4, 93 (2019).

46. Lambrechts, L. Quantitative genetics of Aedes aegypti vector competence for dengue viruses: towards a new paradigm. Trends Parasitol 27, 111–114 (2011).

47. Musso, D., Cao-Lormeau, V. M. & Gubler, D. J. Zika virus: following the path of dengue and chikungunya. Lancet 386, 243–244 (2015).

48. Aliota, M. T. et al. Heterologous Protection against Asian Zika Virus Challenge in Rhesus Macaques. PLoS Negl Trop Dis 10, e0005168 (2016).

49. Lauck, M. et al. Discovery and full genome characterization of two highly divergent simian immunodeficiency viruses infecting black-and-white colobus monkeys (Colobus guereza) in Kibale National Park, Uganda. Retrovirology 10, 107 (2013).

50. Christensen, B. M. & Sutherland, D. R. Exsheathment and Midgut Penetration in Aedes aegypti. Transactions of the American Microscopical Society 103, 423–433 (1984).

51. Rutledge, L. C., Ward, R. A. & Gould, D. J. Studies on the feeding response of mosquitoes to nutritive solutions in a new membrane feeder. Mosqutio News 24, 407–419 (1964).

52. Aliota, M. T. et al. The wMel Strain of Wolbachia Reduces Transmission of Chikungunya Virus in Aedes aegypti. PLoS Negl Trop Dis 10, e0004677 (2016).

